# Diverse Configurations of Erroneous Predictive Coding Across Brain Hierarchies in a Non-Human Primate Model of Autism Spectrum Disorder

**DOI:** 10.1101/2023.11.19.567773

**Authors:** Zenas C. Chao, Misako Komatsu, Madoka Matsumoto, Kazuki Iijima, Keiko Nakagaki, Noritaka Ichinohe

## Abstract

In autism spectrum disorder (ASD), atypical sensory experiences are often associated with irregularities in predictive coding, which proposes that the brain creates hierarchical sensory models via a bidirectional process of predictions and prediction errors. However, it remains unclear how these irregularities manifest across different functional hierarchies in the brain. To address this, we used high-density electrocorticography (ECoG) in a non-human primate model of ASD during an auditory task with two layers of temporal control, and applied a quantitative model to quantify the integrity of predictive coding across two distinct hierarchies. Our results demonstrate that ASD is characterized by sensory hypersensitivity and unstable predictions across two brain hierarchies, and reveal the associated spatio-spectro-temporal neural signatures. Importantly, we observe diverse configurations of underestimation or overestimation of sensory regularities within these hierarchies. This work provides a multi-layered biomarker for ASD, contributing to our understanding of its diverse symptoms.

## Introduction

Autism Spectrum Disorder (ASD) is a neurodevelopmental condition that includes challenges in social interaction and communication, repetitive behaviors, sensory hypo/hypersensitivity, and difficulties adapting to change. Over 250 genes have been identified with strong links to ASD [1], and various brain structures have been associated with autistic traits [2]. However, the neural mechanism of ASD remains elusive, and diagnostic biomarkers are still unavailable [3].

A leading mechanistic investigation of ASD focuses on its atypical sensory perception, such as hypersensitivities to light or sound, which is reported in around 90% of autistic adults [4]. This sensory atypicality has been extensively explained by failures of Bayesian inference [5,6], such as overly-precise sensory observations [7–9], weak prior beliefs [6,10], slow belief updates [11,12], and disbalanced precision controls [10,13,14]. However, the corresponding behavioral evidence are inconsistent and conflicting. For example, prior beliefs in ASD have been shown to be both attenuated [15,16] and intact [8,17,18], and their variability has been reported to be both increased [19] and unaffected [8]. To directly test these Bayesian accounts of ASD, it is critical to identify their underlying neural implementations, which remains unknown.

The most promising implementation of Bayesian inference is predictive coding, which proposes that the brain creates internal models of the sensory world by a hierarchical and bidirectional cascade of large-scale cortical signaling in order to minimize overall prediction errors [20–23]. Specifically, higher-level cortical areas predict inputs from lower-level areas through top-down connections, and prediction-error signals are generated to update the predictions through bottom-up connections when the predicted and actual sensory inputs differ. The theory has been applied to explain how atypical internal models are created in ASD [24,25]. Experimentally, prediction-error signals have been probed by surprise responses when expected stimuli are replaced or omitted. One example is the mismatch negativity (MMN), a late responses to unexpected oddball stimuli in electroencephalography (EEG), which have been reported to show different amplitudes between ASD and typical-developing individuals [26–28]. Different surprise responses in ASD have also been observed in early EEG components [28,29] and blood-oxygen-level-dependent (BOLD) signals [30]. However, meta-analyses on these reports revealed no consistent trend in these differences [31,32].

We hypothesize that the heterogeneous behavioral and neural evidence is caused by a diverse combination of erroneous predictive-coding computations occur across cortical hierarchies, thus cannot be identified by a single neural representation, such as MMN, where prediction-error signals across all hierarchies are mixed together. To test this hypothesis, we extract prediction-error signals across hierarchies and examine their atypical characteristics in ASD. To assess multi-level predictive coding, we use a local-global auditory oddball paradigm, where the subject passively listens to tone sequences with the temporal regularities established at two hierarchical levels [33]. This paradigm allowed a separation of hierarchical prediction-error signals [34–39]. To acquire large-scale neuronal dynamics in ASD with millisecond resolution, we combine high-density hemisphere-wide electrocorticography (ECoG) [40] with a marmoset model of ASD that showed similar functional and molecular features as in human ASD [41]. To provide a mechanistic quantification of erroneous predictive coding, we use a hierarchical predictive-coding model that was previously used to disentangle prediction and prediction-error signals across hierarchies and quantify the integrity of prediction at each hierarchy [42].

Our results reveal sensory hypersensitivity and highly-variable predictions in the autistic animals, which demonstrates the coexistence of the two primary Bayesian accounts of ASD: overly-precise sensory observations and weak prior beliefs. Furthermore, we find distinct patterns of underestimation and/or overestimation of the sensory regularities at different hierarchies in the autistic animals, supporting our hypothesis of erroneous hierarchical predictions as a source of ASD heterogeneity. Our findings map computational theories to their neural implementations and provide a potential neural marker for ASD that is multi-level, high-resolution, and mechanistic.

## Results

### Local-Global Auditory Oddball Paradigm to Establish Hierarchical Regularities

Five marmosets, identified as Ji, Rc, Yo, Ca, and Rm, were used in this study. Among those, Ca and Rm were prenatally exposed to valproic acid (VPA) (see Methods), which was previously used to create a non-human model of ASD [41]. Among 9 VPA-exposed and 10 non-exposed (UE) marmosets in our breeding colony, Ca showed salivary cortisol and diurnal activity levels 1.7 times higher than the average, with a T-score of 69 (n= 19 animals). High cortisol levels and high diurnal activity are characteristic of VPA-exposed marmosets [43], and Ca was considered a model marmoset that is sufficiently exposed to VPA to reproduce ASD. Rm had a twin brother, named Ba, who, according to his breeders, was the least attached to his parents of the more than 200 UE and VPA-exposed marmosets they had raised. Avoidance of parents is a characteristic behavior of ASD. Rm, as Ba, would have been exposed in utero to enough VPA to reproduce ASD.

During the task, subjects were seated with the head fixed and passively listened to a series of short tone sequences based on the local-global auditory oddball paradigm (Figure 1A). Cortical activity was recorded with a 96-channel ECoG array covering nearly an entire cortical hemisphere (left hemisphere for Ji, Ca, and Rm, and right hemisphere for Rc and Yo) (Figure 1B). For Rm, 5 channels in the orbital frontal area and 3 channels in the temporal area were surgical removed during the implantation due to tissue adhesions (88 channels remained). Moreover, the data were collected at different institutes, each with a different data acquisition system, where Ji and Rc from one institute and Yo, Ca, and Rm from the other (see Methods).

**Figure 1.**
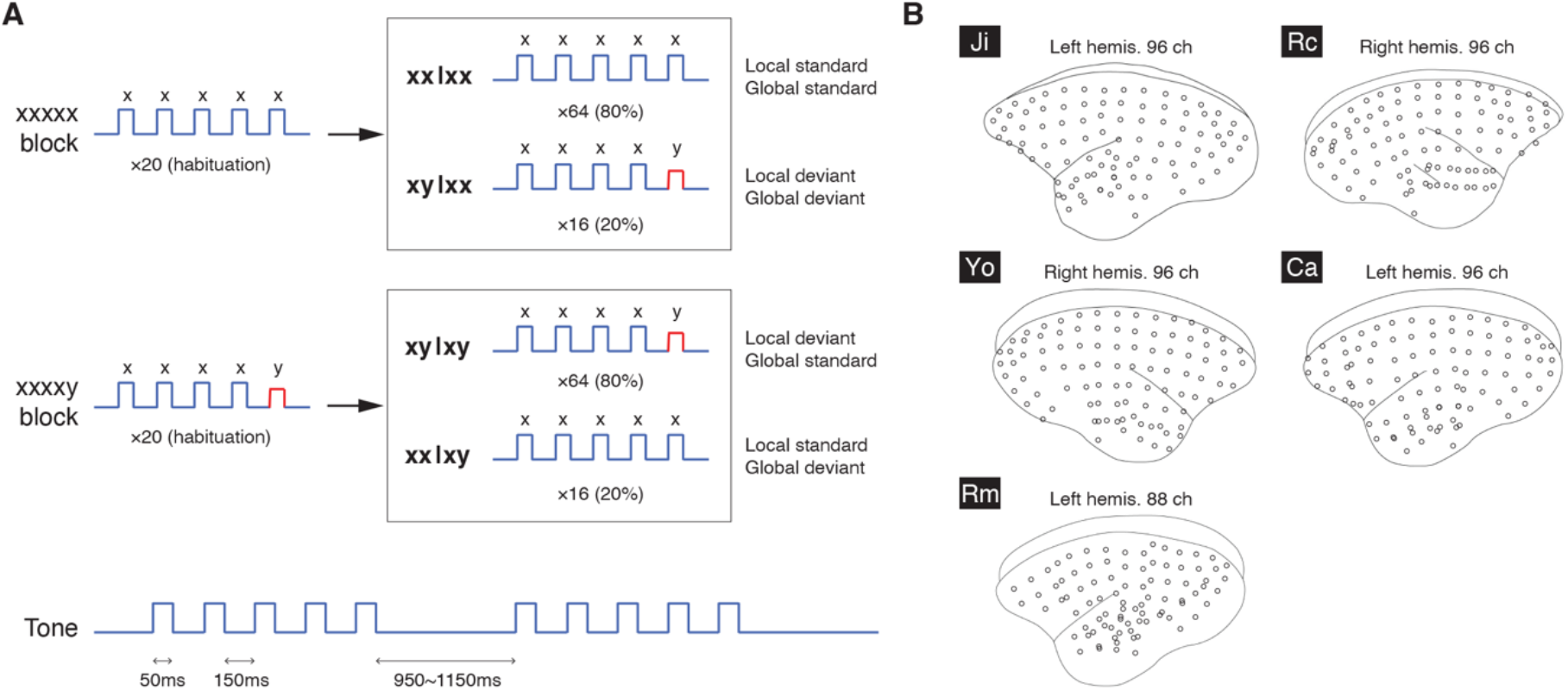
Local-Global Paradigm and ECoG Layouts. (A) The local-global paradigm and the tone and sequence designs. (B) The layout of the 96-channel ECoG arrays in the five subjects. For Rm, 8 channels were surgical removed after implantation due to tissue adhesion, and 88 channels remained.

During each trial, a series of 5 tones were delivered (Figure 1A). The first 4 tones were identical, either low-pitched (tone A) or high-pitched (tone B) (jointly denoted as the standard tone x), and the fifth tone could be either the same (tone x) or different (jointly denoted as the deviant tone y). This resulted in two types of sequences: xx sequence (AAAAA or BBBBB) and xy sequence (AAAAB or BBBBA). Tone sequences were delivered in blocks of 100 trials, where two types of blocks were used: xx or xy blocks. In the xx block, 20 xx sequences were initially delivered as a standard sequence to habituate the subject; then there was a random mixture of 64 xx sequences (the trial type is denoted by xx|xx: xx sequence in xx block) randomly mixed with 16 xy sequences (xy|xx: xy sequence in xx block). Conversely, in the xy block, 20 xy sequences were initially delivered as a standard sequence, followed by a random mixture of 64 xy sequences (xy|xy: xy sequence in xy block) and 16 xx sequences (xx|xy: xx sequence in xy block).

This paradigm was designed to establish two levels of temporal regularity. A local regularity is established within a trial by the repetition of the first 4 tones, which is either followed or violated by the fifth tone. A global regularity is established by habituating the subject to a 5-tone sequence, which is either followed or violated by subsequent sequences. Local and global regularities are orthogonally varied, yielding four trials types: local and global standards (xx|xx), local and global deviants (xy|xx), local deviant but global standard (xy|xy), and local standard but global deviant (xx|xy).

### Deviant Responses to Local and Global Regularity Violations

To examine the effect of VPA on how the local and global regularities were learned and represented in the brain, we evaluated the deviant responses in the brain when the regularities were violated. We compared ECoG signals from the xy and xx sequences in both the xx and xy blocks, i.e. xy|xx – xx|xx and xy|xy – xx|xy. By contrasting xy|xx and xx|xx trials, we can isolate deviant responses that arise when both local and global regularities are violated, i.e. a local deviant response that is also unpredicted by the global rule. Similarly, by contrasting xy|xy and xx|xy trials, we can capture the local deviant response that is predicted by the global rule.

To analyze the large-scale ECoG data, we first identified signal sources over the 96 electrodes (or 88 in Rm) by independent component analysis (ICA) (see Methods). Each independent component (IC) represented a cortical area with statistically-independent source signals (see examples of ICs from Ji in Figure 2A). ICA could identify reference signals (e.g. IC 1 in Figure 2A) and allowed us to bypass re-referencing procedure, such as common averaging re-referencing, which could create substantial bias across all electrodes [Refs]. Furthermore, ICA could help identify spatially-overlapped signal sources (e.g. ICs 59 and 89 in Figure 2A), and extract artifacts (e.g. IC 96 in Figure 2A). Therefore, our further analysis was preformed based on individual ICs, instead of individual channels.

**Figure 2.**
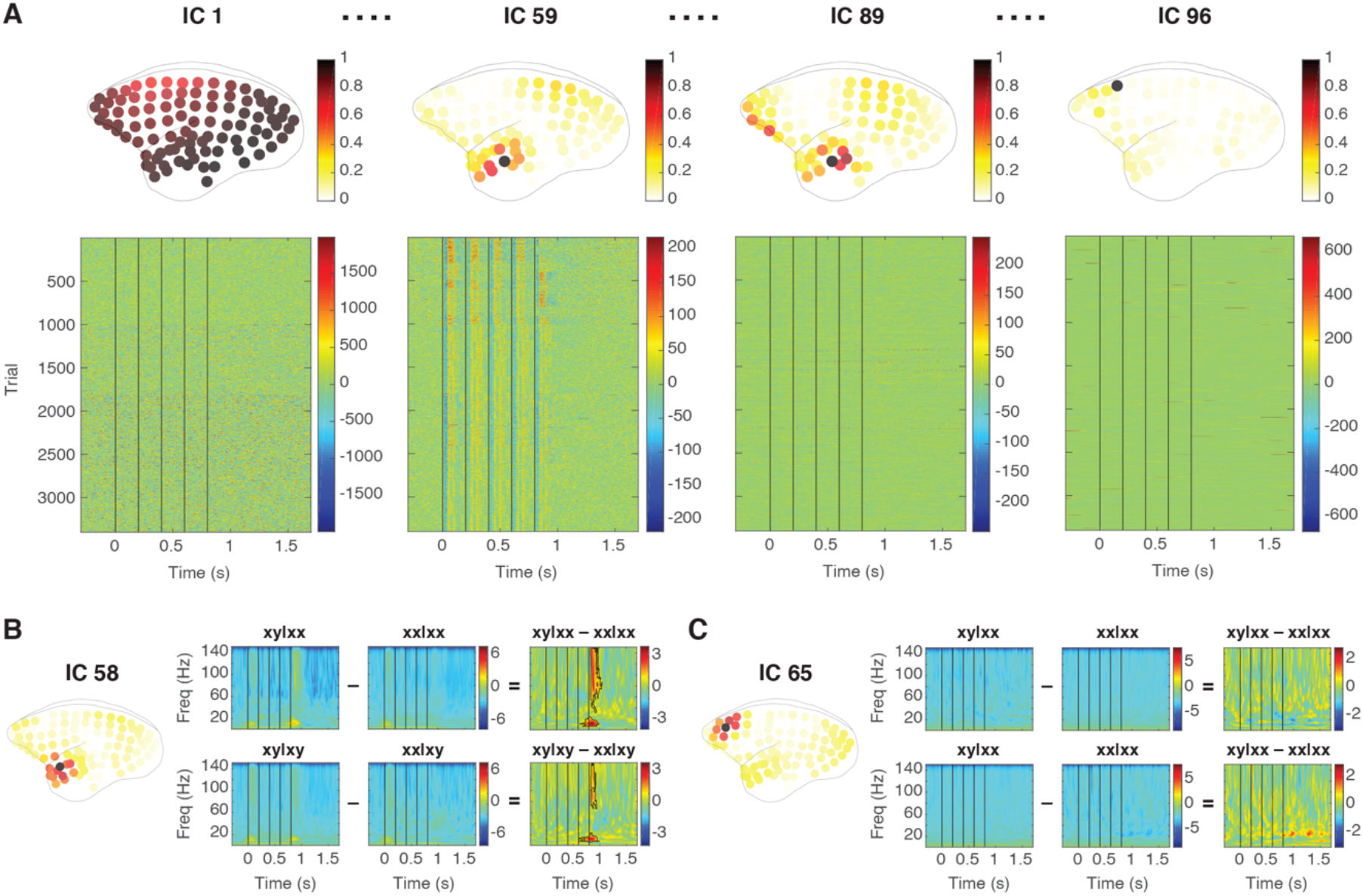
Source Signals and Deviant Responses. (A) Examples of ICs from Ji. For each IC, the absolute spatial contributions were normalized by the maximal value across electrodes, and shown on the top panel. The time courses of all the trials are shown in the bottom panel. Time zero represents the onset of the first tone, and the vertical black lines indicate the onsets of the 5 tones. (B) The deviant responses from IC 58 in Ji. The spatial contribution of the IC is shown on the left. The ERSP for each trial type and the corresponding contrasts are shown. The black contours indicate the deviant response with a significant difference in ERSP in contrasts xy|xx – xx|xx and xy|xy – xx|xy. (C) Example of a non-significant IC. The same representation is used as in panel B.

The spatio-spectro-temporal dynamics of ECoG signals were quantified by the event-related spectral perturbation (ERSP) measured in decibel (dB) (with the baseline from 300 to 0ms before the onset of the first tone, see Methods). Each ERSP represents the in-trial cortical dynamics from an IC, during the time from 300ms before the first tone to 900ms after the fifth tone (a total of 600 time bins), across the frequencies between 0 and 150Hz (a total of 150 frequency bins). Examples of ERSP for all four trial types and their contrasts are shown in Figure 2B for IC 58 (located in the anterior temporal lobe) and in Figure 2C for IC 65 (located in the dorsal prefrontal cortex). A deviant response was defined as a significant difference in ERSP, detected by a nonparametric cluster-based permutation test (α = 0.05 corrected for multiple comparisons, two-sided, see Methods). An IC that showed deviant responses in xy|xx – xx|xx or xy|xy – xx|xy was identified as a *significant IC*. For example, IC 58 was a significant IC with deviant responses in both contrasts (Figure 2B), while IC 65 was not (Figure 2C). The numbers of significant ICs identified in Ji, Rc, Yo, Ca, and Rm were 5, 3, 4, 4, and 5, respectively. All the significant ICs are shown in Figure 3.

**Figure 3.**
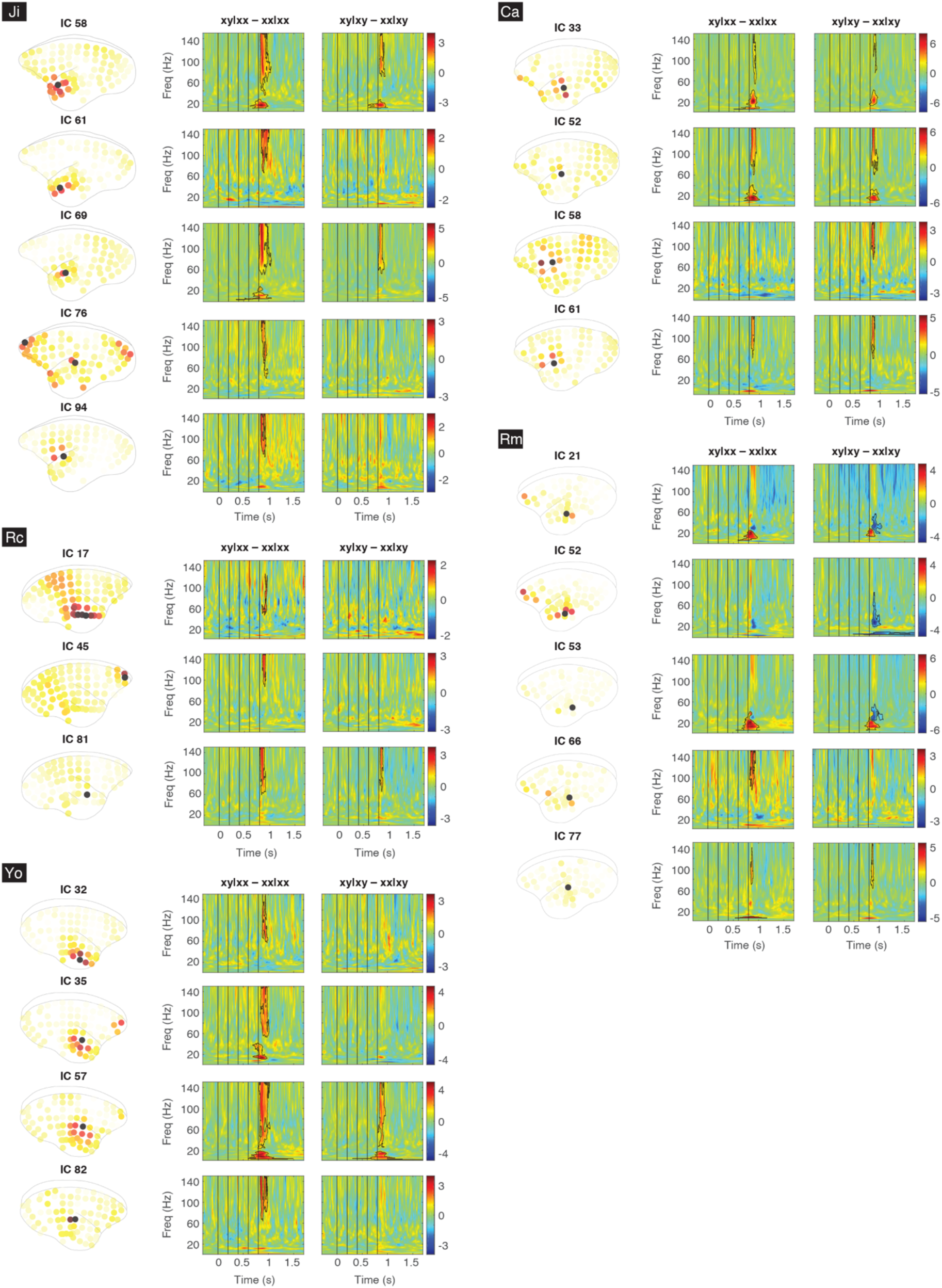
Significant ICs From All Subjects. For each subject, all the significant ICs are labeled and shown with their absolute spatial contributions and the corresponding deviant responses in contrasts xy|xx – xx|xx and xy|xy – xx|xy (black contours).

### Univariate Analysis on Deviant Responses

To compare significant ICs among subjects, we first performed an univariate analysis to quantify their spatial, temporal, and spectral characteristics. To visualize the spatial distribution of each significant IC, its spatial coefficients were normalized to values between 0 and 1 by calculating their absolute values and then dividing them by the maximum (as shown in Figure 3). For each subject, the normalized spatial coefficients were then averaged across all significant ICs to obtain a joint topographic map (Figure 4A), from which we evaluated the relative contributions of three cortical areas: the posterior temporal cortex (pTC), the anterior temporal cortex (aTC), and the anterior prefrontal cortex (aPFC) (Figure 4B). The brain areas were identified based on the Marmoset 3D brain atlas Brain/MINDS NA216 [44] (see Methods). For each area, the relative contribution was quantified by the sum of the spatial distribution in the area divided by the total spatial distribution across all channels. For all subjects except Rm, the relative contributions from strong to weak were pTC > aTC > aPFC. For Rm, the contribution in aPFC was 22.0%, which was 2.6 times stronger the other subjects (8.6 ± 2.0%, n= 4 subjects).

**Figure 4.**
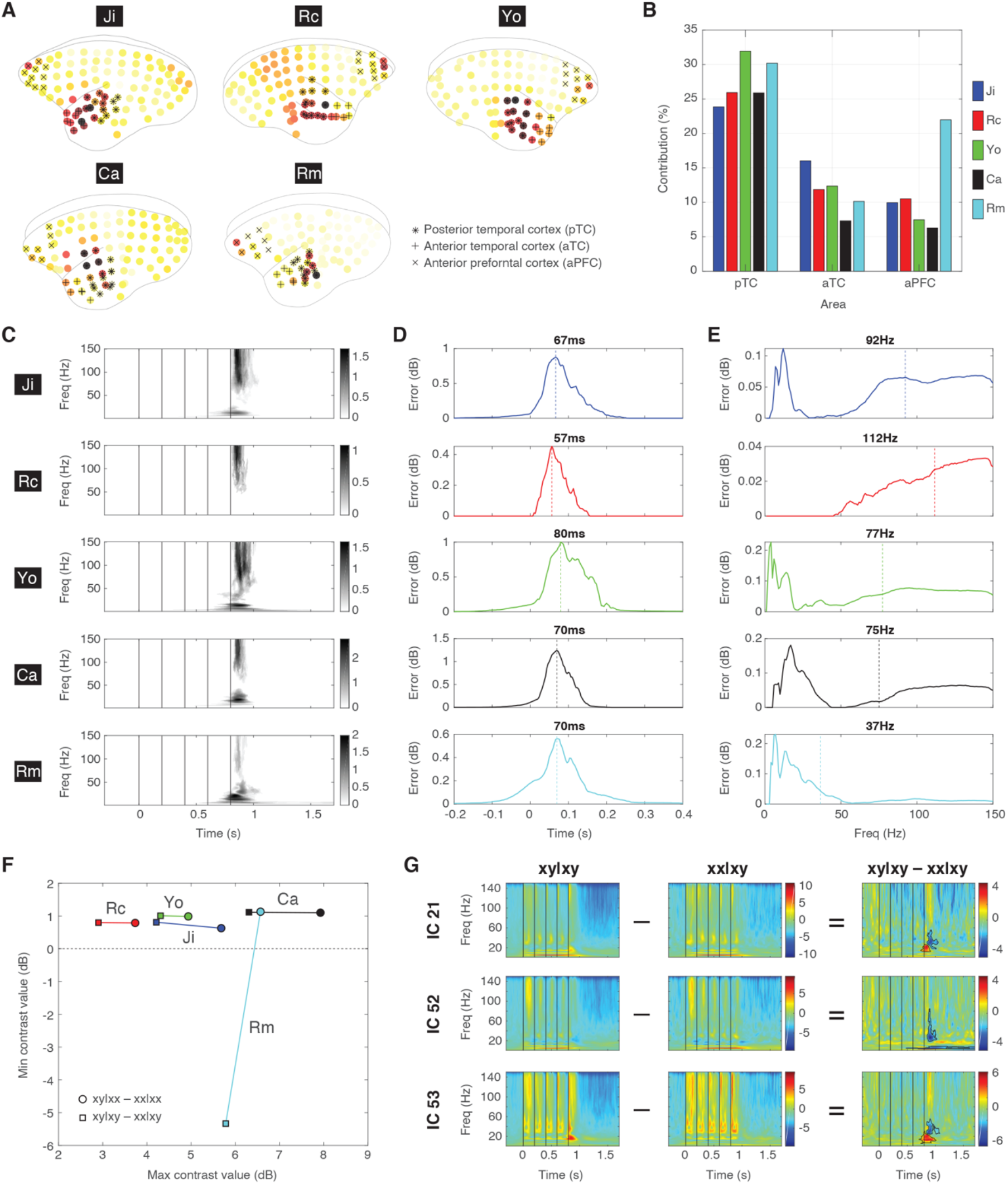
Univariate Analysis of the Significant ICs. (A) The joint topographic map of deviant responses for each subject. The electrodes on pTC, aTC, and aPFC are labeled with black star, plus, and cross signs, respectively. (B) The relative contributions of pTC, aTC, and aPFC for each subject. (C) The joint time-frequency representation of deviant responses for each subject. (D) The temporal profile of deviant responses. The peak response is indicated by a vertical dashed line and the latency is indicated. (E) The spectral profile of deviant responses. The average frequency is indicated by a vertical dashed line and the value is indicated. (F) The maximal and minimal sizes of the deviant responses for each subject. The contrasts xy|xx – xx|xx and xy|xy – xx|xy are indicated by circles and squares, respectively. The color scheme is shared in panels B, D, E, and F. (G) The negative deviant responses in xy|xy – xx|xy in Rm for three significant ICs (21, 52, and 53, as in Figure 3). The ERSP in xy|xy and xx|xy are also shown.

To visualize the temporal and spectral distributions of significant ICs in each subject, absolute values of the deviant responses were averaged across all significant ICs to obtain a joint time-frequency representation (Figure 4C). By averaging the joint deviant response across frequency bins, the peak responses after the last tone were found with comparable latencies of 67, 57, 80, 70, and 80ms for Ji, Rc, Yo, Ca, and Rm, respectively (Figure 4D). By averaging the joint deviant response across time points, the average frequencies were found in the high gamma band at 92, 122, 77, and 75Hz for Ji, Rc, Yo, and Ca, respectively, and in the low gamma band at 37Hz for Rm (Figure 4E).

We further evaluated the size of the deviant responses by measuring the maximal and minimal contrast values in the deviant responses across significant ICs (Figure 4F). For all subjects, the maximal contrast values for contrast xy|xx – xx|xx were positive and greater than the maximal contrast values for xy|xy – xx|xy. This was consistent with the view that a greater surprise was evoked when both local and global regularities were violated (captured by xy|xx – xx|xx), while a smaller surprise was evoked when the local deviant was predicted by the global rule (captured by xy|xy – xx|xy). Furthermore, VPA-exposed Ca and Rm showed stronger deviant responses than the unexposed Ji, Rc, and Yo. On the other hand, the minimal contrasts values were found to be positive, except in Rm where negative deviant responses were found in xy|xy – xx|xy. This is also shown in Figure 3, where a power decrease in the beta/gamma bands (20∼60Hz) was observed in Rm for ICs 21, 52, and 53, particularly in xy|xy – xx|xy. To further examine the ERSP for those ICs, stronger responses to the last x tone in xx|xy were observed (Figure 4G). This indicated a strong surprise toward the global deviant (last tone x), which was not observed in other subjects (e.g. see IC 58 in Ji in Figure 2B), and suggested that Rm was more sensitive to the violation of the global rule.

The univariate analysis revealed some abnormal characteristics in the deviant responses in Ca and Rm. In summary: (1) VPA-exposed Ca and Rm showed stronger deviant responses than the unexposed, suggesting their hypersensitivity to deviant stimuli; (2) hyperactivity in the prefrontal cortex was found in Rm, not Ca, which could link to its hypersensitivity to the global regularity; (3) high-gamma deviant responses, which were thought to represent bottom-up prediction errors, were absent in Rm.

### A Hierarchical Predictive Coding Model for the Local-Global Paradigm

To further investigate how sensory sensitivity and erroneous predictions could lead to the observed abnormal deviant responses, we used a model-fitting analysis based on a quantitative model of hierarchical predictive coding. The quantitative model can explain the brain responses during the local-global paradigm with a goodness-of-fit closed to the optimal data-driven decomposition, and enable a mechanistic evaluations of the underlying sensory sensitivity and prediction strengths at the local and global levels [42].

The model describes the interactions between prediction and prediction-error signals during the last tone of a sequence after both local and global regularities are learned. It consists of three hierarchical levels (Level S, Level 1, and Level 2) and two streams (x stream and y stream). Level S is the sensory level that receives thalamic input, which was a value between 0 and 1, Level 1 learns and encodes the local regularity, which is the tone-to-tone transition probability (TP), and Level 2 learns and encodes the global regularity, which is the sequence probability (SP). The x and y streams process the tone x and y, respectively.

Figure 5A shows the neural operations in the x stream between Levels S and 1. Level S contains a neuronal population (denoted by x_s_) that receives a sensory input (black arrow) and a prediction signal (green arrow) from Level 1, and sends a prediction-error signal (blue arrow) to Level 1. Level 1 contains a neuronal population (x_1_) that receives the prediction-error signal from Level S, and sends a prediction signal to Level S. If we assume that the strengths of the sensory input and the prediction signal are 1 and *P1_x_* (0 :: *P1_x_* :: 1), respectively, then there are two possible situations: (1) if the last tone is x, then the strength of the prediction-error signal is 1– *P1_x_*, (2) if the last tone is not x, then the prediction error is 0 – *P1x* (a negative value) and the strength of the corresponding prediction-error signal is |0 – *P1_x_*|= *P1_x_* (|•| indicates the absolute value). Absolute values are taken because we assume predictions and prediction errors are encoded in neuronal firing rates [45], which can only have non-negative values. Thus, the prediction-error signal received during the last tone at Level 1 in the x stream (denoted as *PE1_x_*) is either 1 – *P1_x_* or *P1_x_*, where the probability of receiving the former is the transition probability from tone x to x (*TP_x_*) and the probability of receiving the latter is 1– *TP_x_* (see the bar graph in Figure 5A).

**Figure 5.**
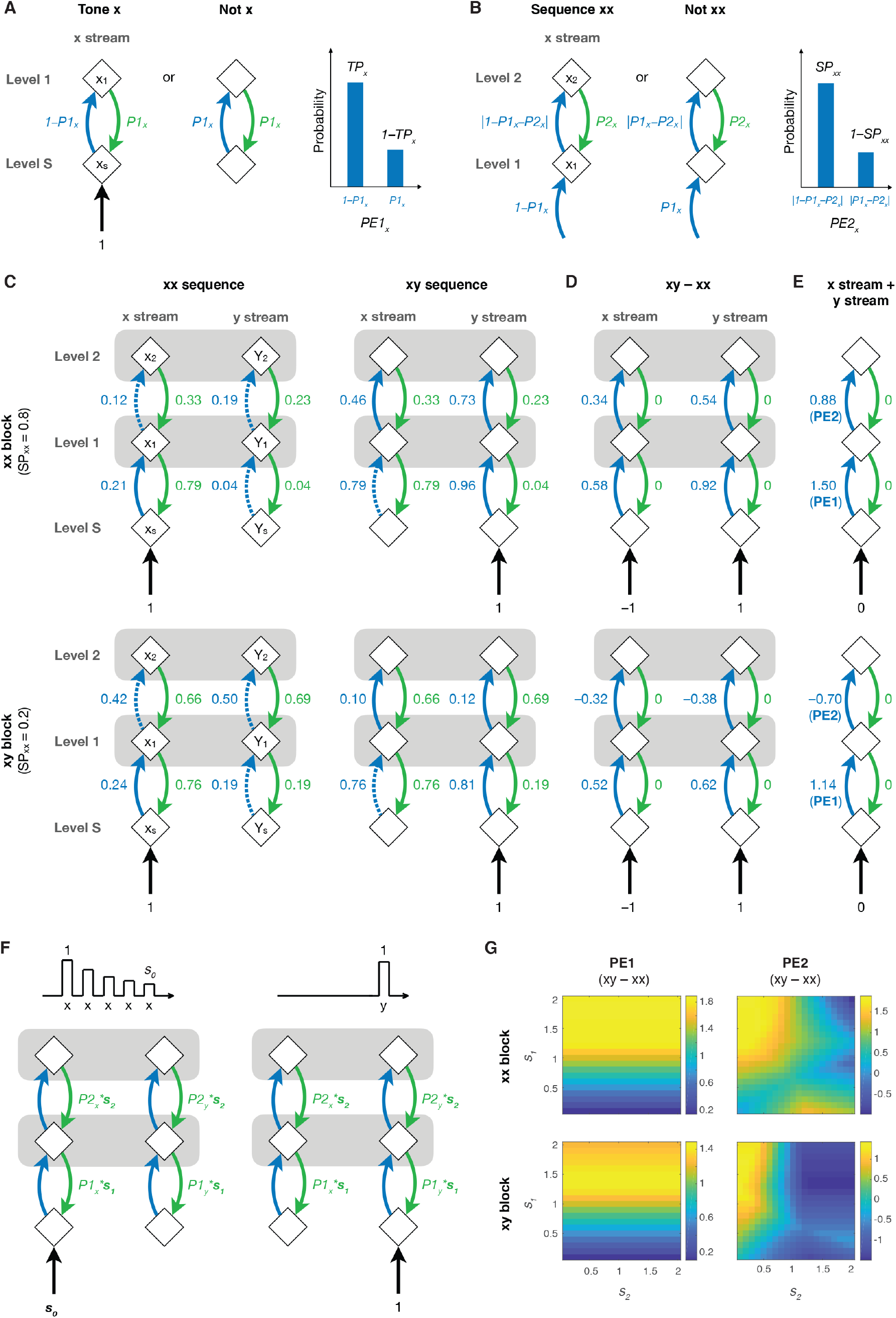
A Quantitative Predictive Coding Model. (A) The proposed neural operations in the x stream between Levels S and 1 during the presentation of tone x or not. An explanatory illustration of the probability distribution of the first-level prediction error in the x stream (*PE1_x_*) is shown on the right. The neuronal populations (diamonds), prediction-error signal (blue arrow), prediction signal (green arrow), and sensory input (black arrow) are shown. (B) The neural operations in the x stream between Levels 1 and 2. An explanatory probability distribution of the second-level prediction error in the x stream (*PE2_x_*) is shown on the right. (C) The complete model during the last tone in xx and xy sequences in the xx and xy blocks. The horizontal gray bars at Levels 1 and 2 indicate integration between the x and y streams for computing transition and sequence probabilities, respectively. The negative errors, where the prediction is greater than the input or prediction error to be predicted, are shown in blue dashed arrows. (D) The contrast values (xy – xx) obtained from panel C. (E) The model values of PE1 and PE2 in the deviant responses by combining contrast values from the x and y streams in panel D. (F) Model tunings with *s_0_*, *s_1_*, and *s_2_*. A decreased response (scaled by *s_0_*) to repeated tone x during the xx sequence (left), and a fresh response to tone y during the xy sequence (right). The corresponding models of the last tone are also shown, where P1 and P2 are scaled by *s_1_*, and *s_2_*, respectively. (G) An example of PE1 and PE2 in the deviant responses in the xx and xy blocks under different combinations of *s_1_* and *s_2_* when *s_0_*= 1.

Figure 5B shows the neural operations in the x stream between Levels 1 and 2. Similar to Level 1, Level 2 contains a neuronal population (x_2_) that receives the prediction-error signal from Level 1, and sends a prediction signal *P2_x_* to Level 1. If the sequence is xx, then the prediction-error signal received at Level 1 is 1 – *P1_x_* (since Level S receives tone x) and the prediction-error signal received at Level 2 is |1 – *P1_x_* – *P2_x_*|. If the sequence is not xx, then the prediction-error signal received at Level 1 is *P1_x_* (since Level S receives not *x*) and the prediction-error signal received at Level 2 is |*P1_x_* – *P2_x_*|. Thus, the prediction-error signal received during the last tone at Level 2 in the x stream (denoted as *PE2_x_*) is either |1 – *P1_x_* – *P2_x_*| or |*P1_x_* – *P2_x_*|, where the probability of receiving the former is the sequence probability of sequence xx (*SP_xx_*) and the probability of receiving the latter is 1– *SP_xx_*.

Based on the model, the strengths of the prediction signals (e.g. *P1_x_* and *P2_x_* in the x stream) are to minimize the mean-squared error received at that level, and can be determined once the transition and sequence probabilities are known (see Methods). Figure 5C shows the complete model for the xx and xy sequences in the xx and xy blocks, which includes the values of prediction and prediction-error signals at both the local and global levels in both the x and y streams. Note that the same prediction signals appear for both the xx and xy sequences, since predictions occur before the last tone arrives. Furthermore, even though the x and y tones are processed in separate streams based on the tonotopic organization, two streams need to integrate information at Levels 1 and 2 to compute transition probabilities (*TP_x_* and *TP_y_*,) and sequence probabilities (*SP_xx_* and *SP_xy_*), respectively. In Figure 5C, we indicate these integrations for probability computations as horizontal gray bars between populations x_1_ and y_1_ and between populations x_2_ and y_2_.

The deviant responses in xy|xx – xx|xx and xy|xy – xx|xy then can be calculated for the x and y streams by subtracting the model values for the xx sequence from those for the xy sequence in each block (Figure 5D). Note that only the prediction-error signals were left in the deviant responses since the same prediction signals are shared between xx and xy sequences. Furthermore, even though x and y streams are modeled separately, their prediction-error values were combined for later model-fitting (Figure 5E). This is based on the assumption that ECoG recordings offer insufficient spatial resolution to separate the x and y streams. As results, the local prediction error (PE1) and global prediction error (PE2) contained the deviant responses are 1.50 and 0.88, respectively in xy|xx – xx|xx, and are 1.14 and –0.70, respectively in xy|xy – xx|xy.

### Models with Erroneous Sensory Sensitivity and Hierarchical Predictions

The model values shown in Figures 5C-5E represent the optimal predictions, where the mean-squared prediction errors are minimized at each level. To further evaluate the potential erroneous sensory sensitivity and predictions for the VPA-exposed, we further added some tunings to the model across different levels (Figure 5F).

At Level S, a scaling factor *s_0_* was added to the sensory input in the x stream to account for the sensory sensitivity or adaptation for the repetitive tone x. The value of *s_0_* was between 0 and 1, where *s_0_=* 1 represents no sensory adaptation or no diminished responses to repeated exposure of tones. For the xy sequence, since tone y does not repeat, adaption does not occur in the y stream. At Levels 1 and 2, we added scaling factors *s_1_* and *s_2_* to the first-level predictions (*P1_x_* and *P1_y_*) and the second-level predictions (*P2_x_* and *P2_y_*), respectively, to account for imperfect predictions. When *s_1_*= 1 and *s_2_*= 1, the predictions are optimal. When *s_1_*< 1 or *s_2_*< 1, the prediction underreacts to the input (sensory input or first-level prediction error, respectively), i.e. “hypo-prediction”, and is insufficient to cancel it out. For example, if *s_1_*= 0, there will be no first level prediction, and the prediction errors continue to propagate to Level 2 without reducing. When *s_1_*> 1 or *s_2_*> 1, the prediction overreacts to the input, i.e. “hyper-prediction”, where the corresponding transition or sequence probabilities are overestimated and additional errors are created. Note that *s_1_* and *s_2_* were applied to both the x and y streams, since erroneous estimation of transition or sequence probabilities could occur at both streams. In Figure 5G, we show an example of PE1 and PE2 contained in the deviant responses in the xx and xy blocks under different combinations of *s_1_* (between 0 and 2) and *s_2_* (between 0 and 2) when *s_0_*= 1 (no sensory adaptation).

### Model-Fitting for Optimal Decomposition of Deviant Responses

We then evaluated which combination of *s_0_*, *s_1_*, and *s_2_* could best explain the observed deviant responses. To achieve this, we first pooled all deviant responses (as shown in Figure 3) to create a tensor with three dimensions: *Contrast*, *IC*, and *Time-Frequency* for the functional, anatomical, dynamical aspects of the data, respectively. For each subject, the dimensionality of the tensor was 2 (xy|xx – xx|xx and xy|xy – xx|xy) by 3∼5 (the number of significant ICs) by 90,000 (600 time points and 150 frequency bins).

We then factorized the 3D tensor into PE1 and PE2 components by performing parallel factor analysis (PARAFAC), a generalization of principal component analysis (PCA) to higher-order arrays [46], with the first dimension fixed with the model values (see Methods). This model-fitting analysis was performed for 9,261 (= 21*21*21) models, each with a unique combination of the scaling factors *s_0_* (21 values between 0 and 1), *s_1_* (21 values between 0 and 2), and *s_2_* (21 values between 0 and 2). For each model, the goodness-of-fit was evaluated by the residual sum of squares (RSS) and core consistency [47]. The best-fitting model was determined as the one with the smallest RSS and a core consistency above 80%.

The parameters of the best-fitting models for all subjects are shown in Figure 6. The best-fitting models for the unexposed (Ji, Rc, and Yo) were found with similar scaling factors: *s_0_* = 0.35∼0.45, *s_1_* = 0.8∼0.9, and *s_2_* = 0.7∼0.8. For VPA-treated, the best-fitting model for Ca was found when *s_0_* = 0.75, *s_1_* =0.3, and *s_2_* = 0.2. This suggested a hyper sensory sensitivity (*s_0_* was twice the size as for the unexposed) and hypo-predictions at both the local and global levels. On the other hand, the best-fitting model for Rm was found when *s_0_* = 0.95, *s_1_* =1.0, and *s_2_* = 1.72. This indicated that Rm shared a similar hyper sensory sensitivity as in Ca, but with a normal local prediction and a hyper global prediction.

**Figure 6.**
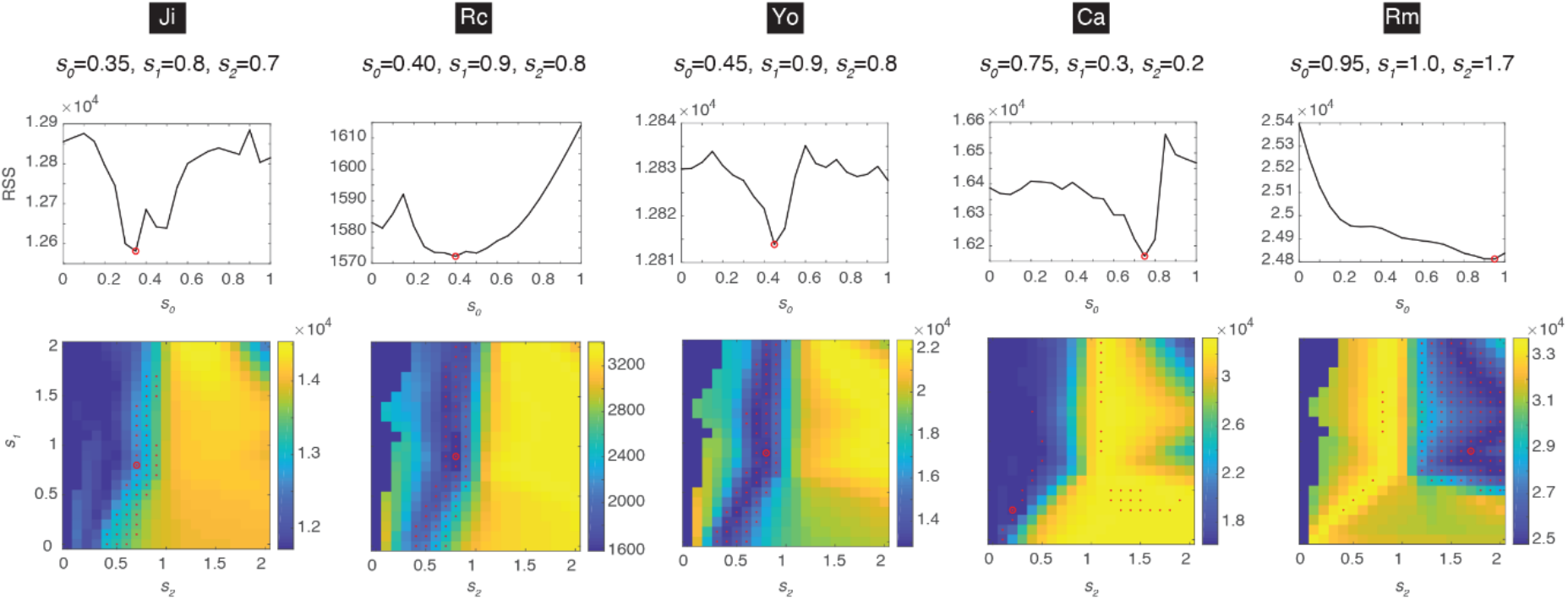
The Optimal Parameter for Model Fitting. The model-fitting results for each subject are shown in each column, where the optimal parameters are indicated. For each *s_0_*, the minimal RSS across different combinations *s_1_* and *s_2_* is shown in the top panel. The minimal RSS is indicated by a red circle. The combination of *s_1_* and *s_2_* under this minimum is indicated by a red circle in the bottom panel. Models with a fitting consistency >80% are indicated by red dots. The color bar represents RSS.

In summary, the model-fitting analysis revealed potential mechanisms that cannot be observed by univariate analysis, and indicated that (1) predictions in the unexposed were close to optimal at both hierarchical levels, (2) hyper sensory sensitivity was found in both VPA-treated, and (3) different types of erroneous hierarchical predictions were observed for VPA-treated.

### Prediction-Error Signals Extracted From Best-Fitting Models

Next we visualized the spatio-spectro-temporal patterns of the PE1 and PE2 components extracted from the best-fitting models. These components were visualized by their composition in the three tensor dimensions. The first dimension showed how much PE1 and PE2 contributed to the deviant responses in the two contrasts (Figure 7A), which was determined by the model and used for the model-fitting. The model values were different across subjects, since different optimal parameters were obtained.

**Figure 7.**
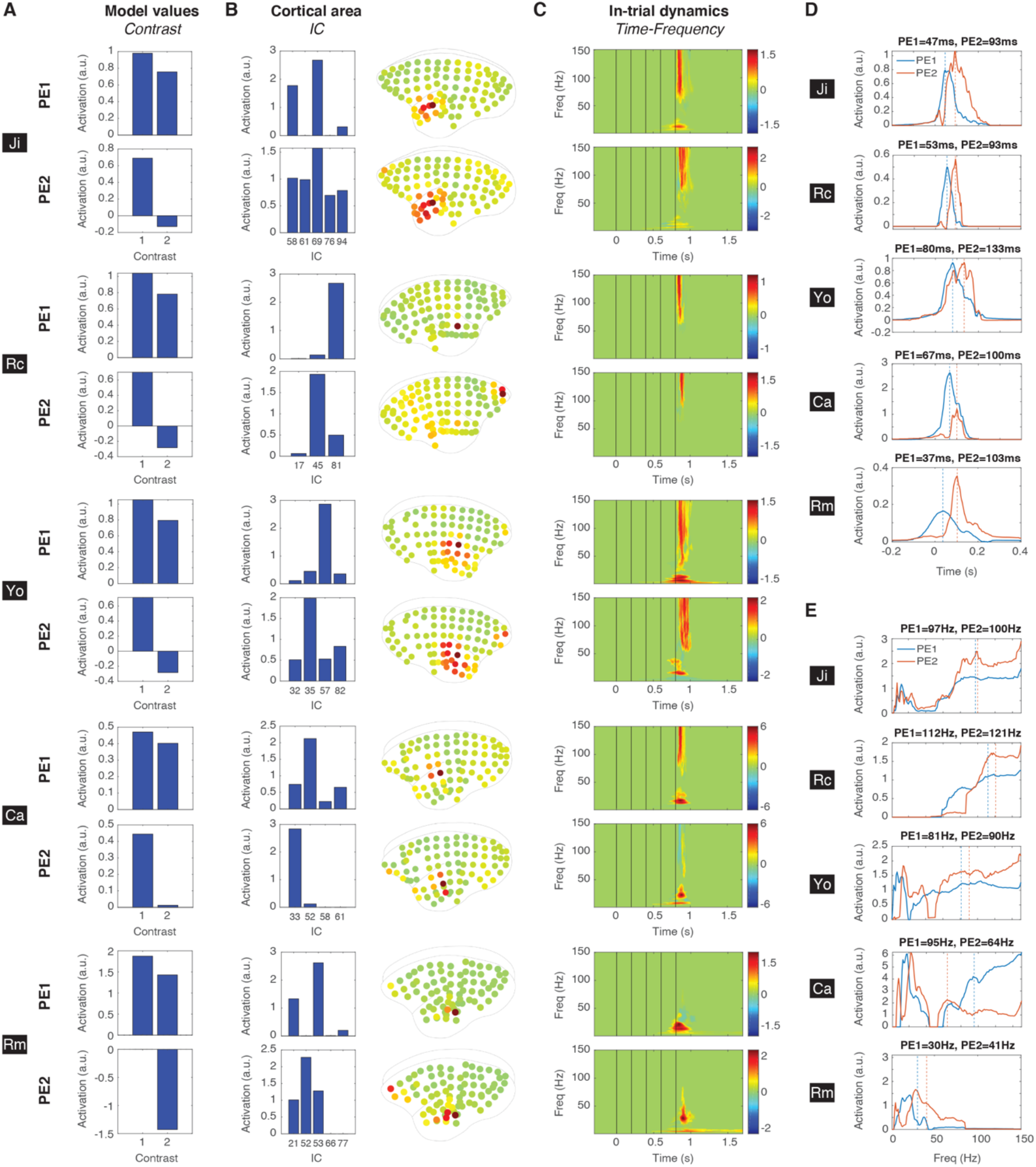
Prediction-Error Signals Extracted From the Best-Fitting Model. (A) The contributions of PE1 and PE2 to the deviant responses in the two contrasts, which are based on the model values from the best-fitting model. (B) The spatial dimension of the PE1 and PE2 components extracted from the best-fitting model. The contribution of each significant IC to PE1 and PE2 and the corresponding average brain maps are shown. (C) The spectro-temporal dimension of the PE1 and PE2 components extracted from the best-fitting model. (D) The temporal profiles of PE1 (blue) and PE2 (orange). The maximal activations are indicated as vertical dashed lines and the corresponding peak latencies are shown. (E) The spectral profiles of PE1 and PE2. The average frequencies activations are indicated as vertical dashed lines and the corresponding values are shown.

The second dimension showed the contribution of each significant IC to PE1 and PE2 (Figure 7B). For example, in Ji, only ICs 58, 69, and 94 contributed to PE1, and IC 69 contributed the most. To further visualize these contributions on a brain map, the normalized spatial distributions of significant ICs (as in Figure 3) were combined based on their contributions (see more details in Methods). The resulting brain maps are shown in Figure 7B. Overlaps between PE1 and PE2 were observed for most subjects, but primarily PE1 appeared in the posterior temporal cortex and PE2 appeared in the anterior temporal cortex and the anterior prefrontal cortex. This propagation of prediction errors from the temporal cortex to the prefrontal cortex is consistent with previous evidence from both monkey and human studies using the local-global paradigm or its variations [33–36,39].

The third dimension showed the in-trial spectro-temporal dynamics for PE1 and PE2 (Figure 7C). To examine the temporal dynamics of PE1 and PE2, we averaged the time-frequency representation in Figure 7C across all frequency bins (Figure 7D). PE1 peaked at 47, 53, 80, 67, and 37msec after the last tone, while PE2 peaked later at 93, 93, 133, 100, and 103msec for Ji, Rc, Yo, Ca, and Rm, respectively. To examine the spectral profiles of the PE1 and PE2 components, we measured their maximal activation at each frequency bin across all time bins (Figure 7E). The average frequencies were 97, 112, 81, 95, and 30Hz for PE1, and 100, 121, 90, 64, and 41Hz for PE2 in Ji, Rc, Yo, Ca, and Rm, respectively. The high-gamma components were absent in Rm, as described in Figure 4E.

### Response Variability Underlying Deviant Responses

For Rm, the absence of high-gamma components in the deviant responses could result from two possibilities: (1) the sizes of prediction-error signals carried in the high-gamma band were comparable between the xx and xy sequences, or (2) the sizes of prediction-error signals were different between the xx and xy sequences but the trial-to-trial variability was too high to obtain statistical significance. The former suggests that no prediction was established, and the latter suggests that the prediction was highly variable over trials.

To test these two possibilities, we measured the trial-by-trial sizes of PE1 and PE2 by evaluating how much PE1 and PE2 contributed to single-trial EEG responses. This was achieved by projecting the single-trial EEG responses onto the spatio-spectro-temporal structures of PE1 and PE2. The spectro-temporal structures of PE1 and PE2 were determined by normalizing the corresponding time-frequency representations (as in Figure 7C) to values between 0 and 1 and then averaged across subjects (Figure 8A). A mask in the high-gamma band was determined as the top 75% values in frequencies above 40Hz (red contour in Figure 8A). The single-trial contributions of PE1 and PE2 were then obtained by projecting single-trial EEG responses of each significant IC onto these masks. For each trial, the projection values were averaged, weighted by the spatial contributions (as in Figure 7B), and resulted in two projection values to describe how much PE1 and PE2 appeared in the high-gamma band (see details in Methods). An example of the projection values is shown in Figure 8B.

**Figure 8.**
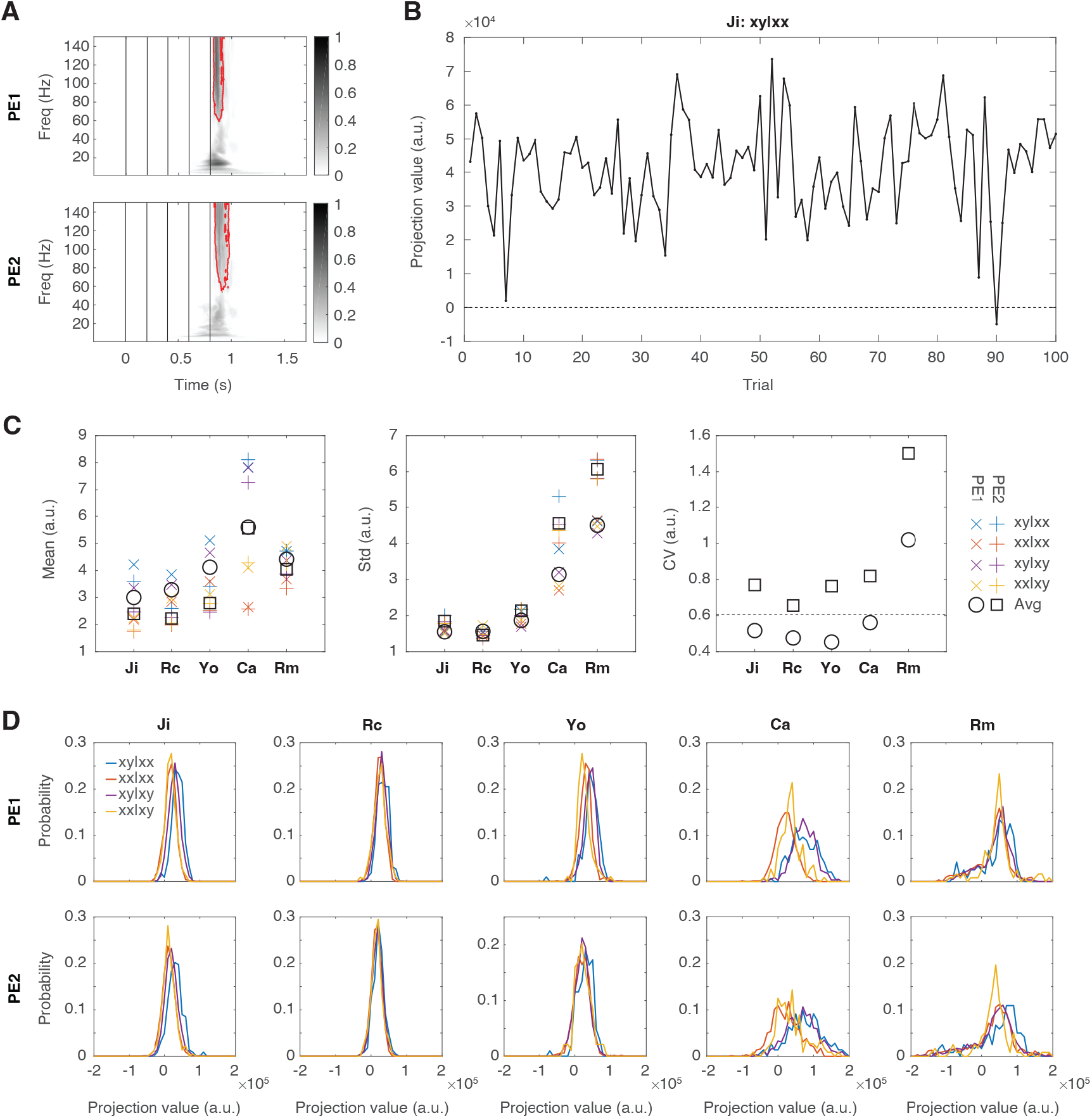
Signal Variability Underlying Absent High-Gamma Components. (A) The spectro-temporal structures of PE1 and PE2. The masks of the top 75% values in the high-gamma band (> 40Hz) are indicated by red contours. (B) An example of the projection values during 100 trials of xy|xx in Ji. (C) The mean, standard deviation, and CV of the single-trial projection values. The mean and standard deviation measured for each trial type are labeled with different colors. The average values across all trial types are indicated by black circles and squares for PE1 and PE2, respectively. CV calculated from the average values are shown. The horizontal dashed line indicates the mean CV from the unexposed (across PE1 and PE2). (D) The probability distributions of the projection values for PE1 (top row) and PE2 (bottom row) in each subject. The probability distribution for each trial types are shown in different colors (same color scheme as in panel C).

For each trial type, the mean, standard deviation, and coefficient of variance (CV, the standard deviation divided by the mean) of the projection values are shown in Figure 8C. The mean activations were higher in Ca compared to the unexposed for both PE1 and PE2, while the means in Rm were comparable to the unexposed. The high mean values in Ca were consistent with the findings that Ca had high sensory sensitivity and the subsequent high prediction errors were not adequately explained away due to hypo-predictions. Furthermore, Ca and Rm showed high standard deviations for both PE1 and PE2. In Rm, the high standard deviation with the comparable mean led to high CV, which supported the second possibility that the absence of the high-gamma components was resulted from highly-variable predictions. In Ca, the high standard deviation was compensated by the high mean activation, which led to low CV and the significant high-gamma components. In addition, PE1 and PE2 were both greater in xy|xx than in xx|xx in all subjects, which was consistent with the view that the xy sequence generated greater local and global prediction errors than the xx sequence in the xx block. To further demonstrate high standard deviations in Ca and Rm, we measured the probability distributions of the projection values for PE1 and PE2 for each trial type and subject (Figure 8D). Wider distributions were found in Ca and Rm, suggesting that the predictions and their subsequent prediction errors in both subjects were highly-variable at both the local and global levels.

## Discussion

We combine a passive auditory paradigm with a quantitative model to extract the neural signatures of hierarchical prediction-error signals, and evaluate the integrity of predictive coding in VPA-exposed animals. Through this approach, we unveil both sensory hypersensitivity and unstable predictions in VPA-exposed animals. Notably, these fluctuating predictions present distinct patterns of underestimation and/or overestimation of hierarchical sensory regularities, potentially contributing to the diverse characteristics of ASD. By linking computational theories with their neural underpinnings, our study lays the groundwork for identifying a comprehensive, multi-tiered, and mechanistic neural marker for ASD.

### Similar neural signatures among macaques, humans, and marmosets

The hierarchical organization of prediction-error signals in the auditory local-global paradigm has been examined in humans and non-human primates [35–39,48]. Utilizing both data-driven and model-driven analyses to decompose predictive-coding signals that are not only interdependent but also spatially and temporally overlapping, we have identified the neural signatures of local and global prediction-error signals in macaque ECoG [34] and human EEG [42]. The spatio-spectro-temporal markers observed in both macaques and humans bear similarity to those depicted in Figure 7, showing prediction-error signals in high frequency band (>30Hz) propagating from the auditory cortex to the frontal cortex with a delay of ∼50msec. This suggests a shared neural organization that facilitates hierarchical predictive coding in both human and non-human primates. Moreover, it indicates the potential applicability of our animal model findings to human patients.

### Multi-level regularity processing in individuals with ASD

The local-global paradigm has been utilized to study atypical perception and emotion processing in ASD. In a study of adults with ASD, a smaller MMN was found in the ASD group than in the typically developing (TD) group [27]. Moreover, both groups demonstrated a reduced MMN when the global rule could be anticipated, such as in the xy block, but this reduction was more pronounced in the TD group than in the ASD group. This implies weaker local and global predictions among individuals with ASD. In a study with children (8∼15 year old) with ASD, no significant differences in MMN were found between ASD and TD groups, suggesting that local prediction error was processed normally [26]. When manipulating the global rule to establish various levels of expectation for local deviants, children with ASD responded differently. There was a decrease in frontocortical responses to sequences that were unexpected, whereas there was an increase in late frontal activation in response to anticipated sequences. These findings suggest that there may be abnormalities in global prediction within the ASD population. In addition, individuals with ASD demonstrated MMN in response to violations of local emotion regularity for both faces and music, but their responses to global emotion regularity violations were absent [49]. These results, derived from a group level analysis, suggest a potential deficiency in global prediction within ASD. They are in alignment with other discoveries of unusual contextual modulations in sensory processing in ASD, observed in both non-social settings [50,51] and social contexts [52–54]. However, our results demonstrate the importance of conducting an individual-level analysis, which would help in identifying the potential diversity in abnormalities in multi-level regularity processing.

### Sensory hypersensitivity and aberrant precision control in ASD

Our findings indicate that the VPA-exposed animals exhibited elevated *s_0_* values, suggesting that their responses to repetitive stimuli were not as significantly reduced compared to the healthy controls. We interpretate this lack of sensory adaption as heightened sensory sensitivity, and use this evidence to support the overly-precise sensory observations account of ASD [7–9]. However, sensory hypersensitivity can also be attributed to imbalanced precision controls, where the brain faces challenges in prioritizing sensory information based on its perceived reliability or precision [10,13,14]. In this case, there is a tendency to assign higher weight to low-level sensory details, potentially intensifying the sensitivity to sensory stimuli. Importantly, these theories are not mutually exclusive and can provide complementary insights into the understanding of sensory processing in ASD. To gain a deeper understanding of their respective contributions, employing a trial-by-trial analysis with Bayesian modeling that incorporates precision parameters could be beneficial [55,56]. Additionally, conducting experiments that effectively control the precision of stimuli can provide valuable insights into the interplay between sensory processing and precision weighting [57].

It is worth noting that our model does not differentiate the origins of this presumed sensory adaption to repetitive stimuli, only its outcome. One possible cause is stimulus-specific adaptation, an inhibitory neuronal mechanism observed in both cortical and subcortical structures [58–60]. Another possible cause is predictive coding itself, where the prediction of transitions between identical tones is learned during repetitions, and the repetitive tones generate less surprise over time. To fully explain the data will require a model that includes the interplay between and stimulus-specific adaptation and predictive coding to describe the neural dynamics during each tone in both cortical and subcortical areas.

### Atypical perceptual learning in ASD

Our findings demonstrate erroneous predictions across different cortical hierarchies in VPA-exposed animals. This implies a deficiency in perceptual learning, which is a key characteristics of the ASD phenotype [61]. This deficiency could stem from abnormalities in synaptic plasticity [62] or learning-related changes in neural connections [63], and lead to slow or irregular belief updates [11,12]. Regrettably, our current model has limitations in assessing the learning process as it only represents the signals once the temporal regularities have already been learned and the errors have been minimized. To understand the dynamic process of prediction updating and error minimization, it is crucial to investigate the trial-by-trial signaling that occurs during the learning process.

A Bayesian model known as the hierarchical Gaussian filtering (HGF) has emerged as a promising candidate for understanding prediction updating and error minimization [64]. This model employs precision-weighted prediction errors [20,22,65] and has been utilized to investigate prediction-error signals in the brain during learning [66–70]. However, its current implementation is limited in terms of hierarchical prediction, despite the term "hierarchical" in its name, which primarily refers to a motor aspect of the model. Nevertheless, there is a high demand for further development and application of hierarchical prediction within this model. Another candidate is dynamic causal modeling (DCM), which is also a Bayesian model that can be used to estimate the coupling among brain regions and the changes in coupling over time and across experimental conditions [71]. This method has been utilized in studying MMN [55,72,73], albeit with the constraint that the brain regions of interest had to be predetermined.

### Non-human primate model of ASD

The phylogenetic closeness between nonhuman primates and humans, particularly in molecular, circuitry, and morphological features of the brain, has made these species increasingly attractive as novel models of psychiatric disorders. Boasting a well-differentiated frontal lobe and intricately stratified hierarchical cortical connectivity, their cerebral cortex closely resembles that found in humans [74]. Marmosets, in particular, offer many advantages as model organisms for the developmental disorder autism. For example, early sexual maturation, high reproductive rates, efficient space utilization due to their compact size, the potential for genetic manipulation, and a complex repertoire of social skills [75].

The VPA-exposed model marmosets utilized in this study show deficits in social tasks that require sophisticated and hierarchical internal models [76,77]. This includes the ability to adjust one’s motivation based on observations of others’ behavior and to evaluate the reciprocity of others. It is important to note that gene expression within the marmoset cortex closely matches the postmortem brains of human ASD, suggesting that it more closely resembles humans than any prior rodent model [41]. Notably, gene expression associated with myelin and inhibitory neurons, which are thought to be important for brain computation, is commonly reduced in both individuals with ASD and the VPA-exposed marmosets. Continued studies of hierarchical predictive coding using the VPA-exposed marmosets may provide important insights into the nature of human ASD. This is especially important given the profound social deficits exhibited in ASD.

In summary, we record large-scale high-resolution neural data in a non-human primate model of ASD and identify different neural signatures underlying different predictive coding accounts of ASD. This research has the potential to contribute to the identification of neural markers specific to different subtypes of ASD and shed light on the impact of prenatal VPA exposure on neurodevelopmental pathways leading to ASD.

## Methods

### Animals

We used five adult common marmosets (Callithrix jacchus; three males and two females, 320–450 g, 22–42 months). Before the ECoG arrays were implanted into the monkeys, they were familiarized with the experimenter and experimental settings. The animals had ad libitum access to food and water throughout the experimental period. Two animals (Ji and Rc), were raised and recorded at RIKEN Center for Brain Science, and the other three animals (Yo, Ca, and Rm), were raised and recorded at the National Center of Neurology and Psychiatry (NCNP). Marmosets were housed in an environment maintained on a 12/12-hour light/dark cycle, and given food (CMS-1, CLEA Japan) and water ad libitum. Temperature was maintained at 27-30°C and humidity at 40-50%.

All procedures of the ECoG study at RIKEN were conducted in accordance with a protocol approved by the RIKEN Ethical Committee. All procedures of the VPA preparation and ECoG study at NCNP were conducted in accordance with NIH guidelines and the "Guide for the Care and Use of Primate Laboratory Animals" published by the National Institute of Neurological Research, National Center of Neurology and Psychiatry, and approved by the Animal Research Committee of NCNP.

### VPA treatment

The method for producing VPA-exposed marmosets was identical to the one detailed in our previous work [41]. In short, serum progesterone levels in the female marmosets were monitored once a week to determine the timing of pregnancy. In addition to the blood progesterone level, pregnancy was further confirmed by palpitations and ultrasound monitoring (Ultrasound Scanning; Xario, Toshiba Medical Systems Corp., Tochigi, Japan). We orally administered 200 mg/kg of sodium valproate (VPA, Sigma–Aldrich, St. Louis, MO, USA) seven times from day 60 to 66 after conception to the mother marmosets. We did not observe obvious malformations or deformities in VPA-exposed marmosets.

### Electrode implants

The whole-hemisphere 96-channel ECoG arrays (Cir-Tech Co. Ltd., Japan) were chronically implanted. We epidurally implanted the array into the right hemisphere for Rc and Yo, and the left hemisphere for Ji, Rm and Ca. Eight electrodes (channels 92∼94) from Rm were cut during the implantation due to tissue adhesions. The surgical procedures for electrode implantation have been previously described in detail [40]. The coordinates of recording electrodes were identified on the basis of the combination of pre-acquired MR images and postoperative computer tomography images using AFNI software [78] (http://afni.nimh.nih.gov). Then, we estimated the location of each electrode on cortical areas by registering to the Marmoset 3D brain atlas Brain/MINDS NA216 [44] with AFNI and ANTS [79].

### Experimental setup

ECoG signals from monkeys Ji and Rc were acquired at RIKEN using a Grapevine NIP system (Ripple Neuro, Salt Lake City, UT) at a sampling rate of 1 kHz. Experiments of monkeys Yo, Rm, and Ca were conducted at NCNP. The neural signals were stored at a 1017.25 Hz sampling resolution into a TDT signal processing system RZ2 (Tucker-Davis Technologies, Alachua, FL). During the ECoG recordings, the marmoset was seated in a primate chair in an electrically shielded and sound-attenuated chamber with their head fixed. The auditory stimuli were delivered bilaterally by two audio speakers (Fostex, Japan) at a distance of ∼20 cm from the head at an average intensity of 65 dB SPL.

### Stimuli and experimental procedure

Two tones with different pitches (Tone A = 800Hz; Tone B = 1600Hz) were synthesized. Each tone was 50 ms in duration. Series of five tones were presented with a 150 ms inter-tone interval, with 950-1150 ms was set between the offset of the last tone of a sequence and the onset of the first tone of the following sequence (see Figure 1A). Four different stimulus blocks were used: AAAAA, BBBBB, AAAAB, and BBBBA blocks. In AAAAA blocks, 20 AAAAA sequences were delivered, followed by a random mixture of 64 AAAAA and 16 AAAAB. In BBBBB blocks, 20 BBBBB sequences were delivered, followed by a random mixture of 64 BBBBB and 16 BBBBA. In AAAAB blocks, 20 AAAAB sequences were delivered, followed by a random mixture of 64 AAAAB and 16 AAAAA. In BBBBA blocks, 20 BBBBA sequences were delivered, followed by a random mixture of 64 BBBBA and 16 BBBBB. In each experimental day, we conducted ECoG recordings on 1∼8 blocks, depending on the animal’s condition. For each animal, we performed 7-9 recordings for each block. The ECoG data can be freely downloaded (https://dataportal.brainminds.jp/).

### Data analysis

#### Preprocessing and Independent Component Analysis (ICA)

The ECoG signals were downsampled to 300Hz by EEGLAB on MATLAB [80] (function: pop_resample.m). Bad channels were then removed by visual inspections: channels 6, 7, 8, and 80 were removed in Rc; channels 8, 68, 71, 81, 83, 85, 87, and 88 were removed from Yo; channels 2, 44, 48, 61, 63, 64 were removed in Ca, channels 1, 6, 7, 8, 9, 10, 43, 43, 49, and 81 were removed in Rm. For each subject, all data were concatenated together, and ICA was performed by the FieldTrip Toolbox [81] (function: ft_componentanalysis.m with the runica algorithm). For each trial, the ICA signals were aligned at the onset of the first tone, and signals from 0.3s before to 1.7s after the onset of the first tone were segmented and used for the further analyses.

#### Event-related spectral perturbation (ERSP)

For each subject, independent component (IC), and trial, the time–frequency representation of the ICA signal was generated by Morlet wavelet transformation at 150 different center frequencies (1∼150Hz) with the half-length of the Morlet analyzing wavelet set at the coarsest scale of 7 samples, which is implemented in the FieldTrip Toolbox (ft_freqanalysis.m). Baseline normalization was then performed to calculate the decibel values by using the baseline period from –0.3 to 0s (time zero as the onset of the first tone) (ft_freqbaseline.m).

#### Deviant response

For each subject, the deviant responses (xy|xx – xx|xx and xy|xy – xx|xy) were calculated for each IC across all trials. To measure the significance of the difference in ERSP (as the black contours shown in Figures 2B), we performed permutations by shuffling trial indices, and used a nonparametric cluster-based method for multiple comparisons correction [82], which is implemented in FieldTrip Toolbox (ft_freqstatistics.m with 500 permutations). Non-significant values in the deviant responses were set to 0, and ICs with no significant deviant responses in both xy|xx – xx|xx and xy|xy – xx|xy were considered as non-significant ICs.

#### Model-fitting with parallel factor analysis (PARAFAC)

We used PARAFAC, a generalization of principal component analysis (PCA) to higher-order arrays [46], which was previous used for the computational extraction of latent structures in functional network dynamics [34,83–85]. To decompose deviant responses into components with theorized contrast values, PARAFAC was performed with the first dimension *Contrast* fixed with the values proposed by the model. This was done by the N-way toolbox [86], with no constraint on all three dimensions (using FixMode and OldLoad inputs in parafac.m). The convergence criterion (i.e., the relative change in fit for which the algorithm stops) was set to 1e−6. The initialization method was set to be direct trilinear decomposition (DTLD), which was considered the most accurate method [87]. For each fitting, the residual sum of squares (RSS) and the core consistency diagnostic [47] were measured.

#### Brain spatial contribution

For each significant IC, the absolute values of the spatial filter (1 × number of channels) were first calculated and normalized by their maximal value (as in Figure 3). The brain map shown in Figure 7B was the linear combination of the normalized spatial filters of all significant ICs and their contributions in the model-fitting (Figure 7B).

#### Single-trial projection

For each subject, a single-trial ERSP response (ERSP= number significant ICs × 150 frequency bins × 600 time points) was projected on a PE1 or PE2 spectro-temporal mask (FT= 600 time points × 150 frequency bins) (as in Figure 8A) and the contributions of all significant ICs (S = 1 × number significant ICs) (as in Figure 7B). This was calculated as S*ERSP*FT, which yields a single scalar value. Note that the spectro-temporal masks for PE1 and PE2 were calculated from all subjects and thus shared across subjects, while the contributions of all significant ICs were different across subjects.

### Model calculation

We used a simple model we previously proposed [42]. In the model, the optimal value of each prediction signal is to minimize the mean-squared error received. For example, the mean squares of *PE1_x_* (denoted by *MSPE1_x_*) can be devised as (based on the bar graph in Figures 5A):

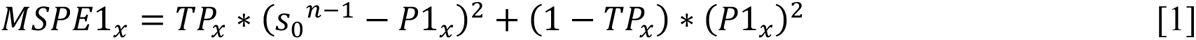

The minimums occur when:

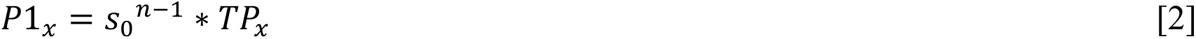

And *P1_y_* can be obtained in the same fashion:

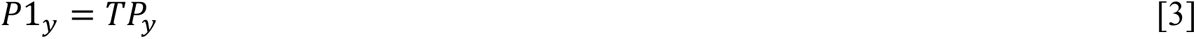

This represent the optimal prediction where first level prediction errors are minimized. Then we added the scaling factor *s_1_* to *P1_x_* and *P1_y_* and calculate the mean squares of *PE2_x_* (denoted by *MSPE2_x_*):

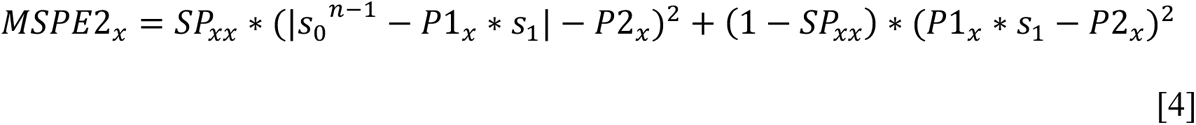

The minimums occur when:

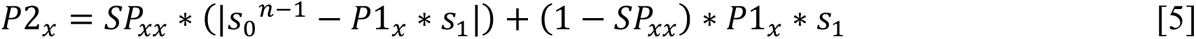

And *P2_y_* can be obtained in the same fashion:

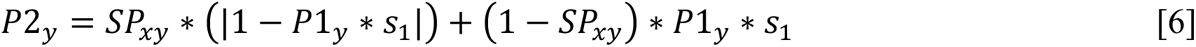

Note that the *P2_x_* and *P2_y_* here represent the optimal predictions when potential erroneous predictions at the first level are considered. Also, *s_2_* was applied to calculate the second level prediction errors, i.e. *P2_x_*\**s_2_* and *P2_y_*\**s_2_* were used (as shown in Figure 5F).

Based on the model, all prediction signals are determined once the transition probabilities (*TP_x_* and *TP_y_*), sequence probabilities (*SP_xx_* and *SP_xy_*), and scaling factors (*s_o_*, *s_1_*, and *s_2_*) are known. The transition probabilities can be calculated based on the number of tones in a sequence and the sequence probabilities (the MATLAB code for these calculations is provided).

## Acknowledgements

We thank Yuri Shinomoto and Takaaki Kaneko for animal care and awake recordings; Junichi Hata for obtaining the MRI images. We also thank Dr. Wataru Suzuki for his assistance in setting up the ECoG lab system and marmoset experiments and Ms. Akiko Tsuchiya for her technical support to marmoset breeding and the creation of the VPA-exposed marmoset. This work was supported by World Premier International Research Center Initiative (WPI), MEXT, Japan (to Z.C.C.), Brain/MINDS from the Japan Agency for Medical Research and Development (JP20dm0207001 and JP20dm0207069 to M.K.), JSPS KAKENHI (JP 23H04978 to M.K.), JST Moonshot R&D (JPMJMS2294 to M.M), and an Intramural Research Grant (Nos. 23-7, 26-9, and 29-6 to N.I.) for Neurological and Psychiatric Disorders from the NCNP and AMED (JP23dm0207066 to N.I.).

## Author contributions

Z.C.C. conceptualized the study. M.K. and Z.C.C. refined the experimental protocol. M.K. coordinated and conducted ECoG experiments. K.I. and M.M. conducted ECoG experiments at NCNP. N.I. and K.N. provided VPA marmosets, including marmoset rearing. Z.C.C. designed and performed the data analysis. Z.C.C. wrote the first draft of the paper, and N.I. and M.K. helped with the editing. All authors contributed to and have approved the final paper.

